# Drosophila activins adapt gut size to food intake and promote regenerative growth

**DOI:** 10.1101/2023.01.26.525639

**Authors:** Christian F. Christensen, Quentin Laurichesse, Rihab Loudhaief, Julien Colombani, Ditte S. Andersen

## Abstract

Rapidly renewable tissues adapt different strategies to cope with environmental insults. While tissue repair is associated with increased ISC proliferation and accelerated tissue turnover rates, reduced calorie intake triggers a homeostasis-breaking process causing adaptive resizing of the gut. Here we show that activins are key drivers of both adaptive and regenerative growth. Activin-β (Act-β) is produced by progenitor cells in response to intestinal infections and stimulates ISC proliferation and turnover rates to promote tissue repair. Dawdle (Daw), a divergent Drosophila activin, signals through its receptor, Babo^C^, in progenitor cells to promote their maturation into enterocytes (ECs). Daw is dynamically regulated during starvation-refeeding cycles, where it couples nutrient intake with progenitor maturation and adaptive resizing of the gut. Our results highlight an activin-dependent mechanism coupling nutrient intake with progenitor-to-EC maturation to promote adaptive resizing of the gut and further establish activins as key regulators of adult tissue plasticity.

## Introduction

Adult tissues with high turnover rates depend on stem cells (SCs) to provide a continuous source of differentiated cells to maintain tissue homeostasis. Adaptation of organ size and physiology to internal or environmental cues involves adjustments of divisions and fate decisions to maintain tissue integrity and ensure organism health and reproductivity. Due to a high level of conservation of both structure, physiology, and the gene-regulatory network controlling ISC activity, the Drosophila midgut has proven a powerful model for dissecting the molecular mechanisms underlying adult SC activity and fate decisions (Liu, Hodgson, and Buchon 2017; Colombani and Andersen 2020). Drosophila ISCs are embedded basally throughout the gut epithelium and divide non-symmetrically to give rise to a daughter ISCs and transient progenitor cells, enteroblasts (EBs) or EE progenitor cells (EEPs), that are destined to differentiate into ECs or EE cells (EECs), respectively (Biteau and Jasper 2014; Guo and Ohlstein 2015; Micchelli and Perrimon 2006; Ohlstein and Spradling 2006; Zeng and Hou 2015). In homeostatic conditions, the majority of ISCs undergo asymmetric divisions giving rise to one ISC (self-renewal) and one daughter cell committed to differentiate into an EB (90%) or pre-EEC (10%) ensuring a constant pool of progenitor cells (de Navascues et al. 2012; Ohlstein and Spradling 2006). By contrast, adaptive growth, which can be triggered by anticipatory internal signals, e.g., following mating (Reiff et al. 2015), or environmental cues, is characterized by accelerated ISC division rates and a shift from asymmetric to symmetric ISC divisions (O’Brien et al. 2011). One striking example of adaptive growth is the shrinkage and expansion of the gut in response to cycles of starvation and refeeding, which has been observed in a wide range of organisms (Altmann 1972; Andrew et al. 2015; Dunel-Erb et al. 2001; Lucchetta and Ohlstein 2017; McLeod et al. 2010; O’Brien et al. 2011; Secor, Stein, and Diamond 1994). Intestinal infections also trigger a switch in division mode towards symmetric divisions to ensure rapid replacement of lost ECs (Hu and Jasper 2019; Tian, Wang, and Jiang 2017; Zhai, Boquete, and Lemaitre 2017). To identify niche-derived signals controlling ISC division rates and/or fate decisions in conditions associated with accelerated tissue turnover rates, we used RNAis to knock down all secreted peptides (approx. 800) in different populations of the ISC niche and screened for reduced survival to enteric infection. Strikingly, among the top candidate hits, we identified two ligands belonging to the activin branch of the TGF-β superfamily, Act-β and Daw, and their inhibitor, Follistatin (FS), suggesting a key role of activin signaling in controlling ISC activity and fate decisions.

The architecture of the Activin signaling pathway is highly conserved between flies and mammals (Song et al. 2018). In mammals, activins and Nodal share the same receptors and effectors, and hence, the pathway is often referred to as the Nodal/Activin pathway. Binding of Nodal/Activin growth factors to two type II activin receptors (ActRIIA/B) results in the recruitment, phosphorylation, and activation of two type I activin receptors (ALK4 or ALK7), which subsequently triggers the phosphorylation of receptor-regulated R-Smads (Smad 2 and Smad 3). Phosphorylated R-Smads associate with the Co-Smad, Smad 4, and translocate into the nucleus to regulate the expression of broad range of genes in a tissue specific and context dependent manner (Pauklin and Vallier 2015). In much the same way, binding of Drosophila activins to a single type I receptor, Baboon, promotes the assembly of heteromeric type I/type II (Put/Wit) receptor complexes triggering phosphorylation of the downstream activin-specific R-Smad-related protein, Smad on X (Smox), which together with the Co-Smad, Medea, translocates to the nucleus to regulate gene expression (Song et al. 2018; Brummel et al. 1999). Alternative splicing of Babo gives rise to three isoforms, Babo^A^, Babo^B^, and Babo^C^, that differ in their extracellular domain, and hence their affinity for the three related activin ligands, Act-β, Daw, and Myostatin (Myo) (Brummel et al. 1999; Jensen et al. 2009). Daw, which is a divergent paralogue of Act-β more related to vertebrate TGFβ1, signals exclusively through Babo^C^ *in vitro* (Jensen et al. 2009), whereas both Myo and Act-β was reported to utilize Babo^A^ (Awasaki et al. 2011; Song et al. 2017; Upadhyay et al. 2020). For now, a clear relationship between BaboB and one or more activins has not been established. Activin signaling is further regulated by the highly conserved secreted antagonist FS, which inhibits ligand/receptor interactions in both flies and mammals.

While role of activin signaling in progenitor expansion and cell specification during early development and organogenesis is well established in both flies and mammals (Brummel et al. 1999; Ellis et al. 2010; Lengil, Gancz, and Gilboa 2015; Ng 2008; Parker et al. 2006; Peterson and O’Connor 2013; Rossi and Desplan 2020; Wells et al. 2017; Yu et al. 2013; Zheng et al. 2006; Zhu et al. 2008; Pauklin and Vallier 2015; Ting et al. 2014), its function in adult tissues is not well understood. For instance, Activin B/ActRIB signaling promotes stemness of hair follicle SCs in the skin (Kadaja et al. 2014), while neutralization of Activin A by the activin antagonist, FS, is required to maintain the proliferative potential of ΔNp63^+^ stem- and progenitor cells in mouse salivary glands (Min et al. 2020). In flies, FS is required in testicular somatic cyst SCs (CySCs) to maintain niche quiescence, as loss of FS in CySCs or ectopic activation of activin signaling in hub (niche) cells results in their transdifferentiation into CySCs and a loss of niche capacity (Herrera et al. 2021). It is not clear whether the opposing outcomes of activin signaling on SCs self-renewal and differentiation reflect tissue specific requirements. Nevertheless, mutations affecting Activin/Nodal signaling are frequent in cancer stem cells (CSCs) in a variety of tissues (Lonardo et al. 2011; Massague 2008; Topczewska et al. 2006), suggesting a broader role of activins in regulating SC fate decisions in adult tissues.

Here we show that activin signalling is instrumental in coordinating ISC proliferation and fate decisions in both homeostatic conditions and during regenerative/adaptive growth. While Act-β signals through BaboA in ISCs to promote their self-renewal and differentiation into EBs, Daw/Babo^C^ signaling is required in EBs for their maturation into ECs. Act-β is produced by EBs in response to intestinal infections and required for tissue repair, while Daw expression is dynamically regulated by nutrient intake and plays a key role in the adaptive resizing of the gut during cycles of starvation and refeeding by controlling EB maturation and tissue turnover rates.

## Results

### Activin signalling regulates homeostatic gut epithelial turnover rates

While activins have essential functions during early development in mammals, their function in most adult tissue including the gut has not been studied. To investigate how activin signalling might regulate tissue turnover in the adult gut, we analysed the expression pattern of the activin type I receptor Baboon (Babo) using a Babo>UAS-GFP reporter (Fig. 1A-A’’). We found that Babo is expressed at high levels in ISCs and EBs (Fig. 1A) and at variable levels in ECs (Fig. 1A), while its expression was not detected in EECs (Fig. 1A). To analyse whether activin signalling is required to sustain tissue turnover in homeostatic conditions, we knocked down Babo and its downstream effector Smox in progenitor cells and monitored the effect on gut size and numbers of newly generated ECs using a Repressible Dual Differential stability cell Marker (ReDDM) line (Antonello et al. 2015). The ReDDM system labels progenitor cells (ISCs/EBs) with long-lived Histone (His) 2B-RFP and a short-lived GFP, while newly generated ECs are only positive for His-2B-RFP that remains stable for 28 days after its expression is turned off in ECs (Fig. 1C). Knockout of either Babo or Smox reduced gut size (Fig. 1B-B’) and slowed down the production of ECs, demonstrating that activin signalling is required in progenitor cells to maintain tissue turnover rates in homeostatic conditions (Fig. 1D-F, H-H’). Consistent with this, ectopic expression of a constitutively active form of Smox in progenitor cells was sufficient to trigger ISC proliferation and accelerate tissue turnover (Fig. 1D, G, H’’-H’’’).

**Figure 1:**
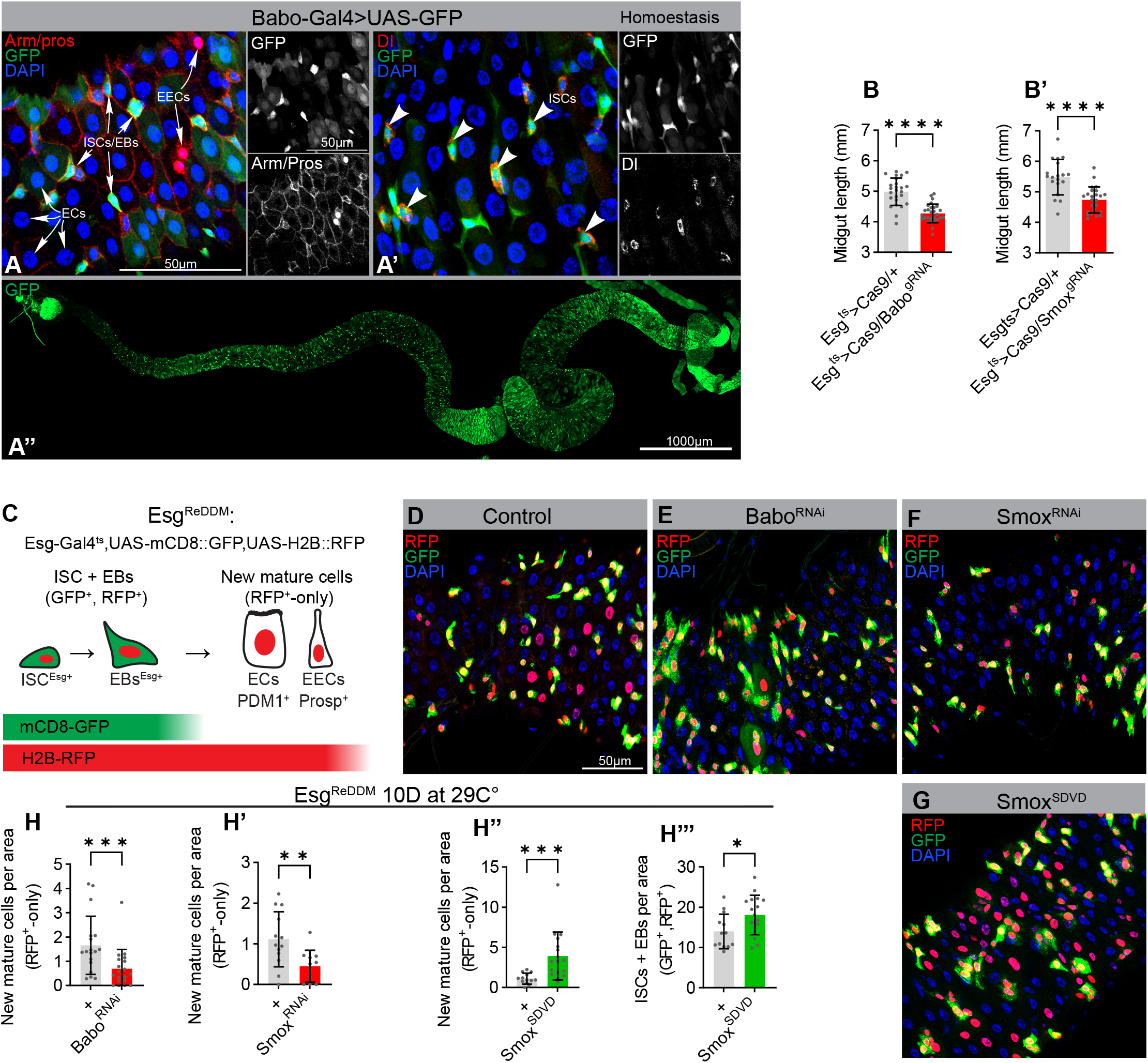
The activin signaling pathway control turnover of intestinal cells. (A-A’’) The Type I Activin receptor Babo is expressed in the midgut. (A) Confocal image of a posterior (A-A’) or whole (A’’) midgut from a Babo-Gal4>UAS-GFP reporter line stained for GFP (green, A-A’’), Armadillo (Arm) and Prospero (Pros) (Red, A) or Dl (Red, A’), and DNA (blue, A-A’) reveal enriched expression in delta-positive ISCs and other diploid cells (EBs) (A’), not marked by nuclear Pros (A). (B-B’) Activin signaling maintains homeostatic gut size. Quantifications of midgut lengths after *escorgot*-Gal4 (esg>)-mediated knockout of Babo or Smox in stem and progenitor cells using CRISPR/Cas9. (C) Schematic diagram of the Esg^ReDDM^ system. Membrane tethered CD8::GFP and nuclear localized H2B::RFP is specifically coexpressed in stem and progenitor cells with the *esg*> driver. As progenitors differentiate into mature cells, loss of *esg*-Gal4 expression terminates further production of GFP and RFP. Differential stability of the fluorophores results in rapid degradation of GFP whereas RFP remains in differentiated cells for an extended duration of time. (D-H’’’) Activin signaling maintains homeostatic cell turnover. Representative confocal images of dissected posterior midguts from control flies (D) and flies expressing Babo^RNAi^ (E), Smox^RNAi^ (F) and Smox^SDVD^ (G) in stem and progenitor cells using Esg^ReDDM^ tracing for 10 days at homeostatic conditions and quantification of cell turnover in each condition (H-H’’’). Expression of Babo^RNAi^ and Smox^RNAi^ reduces the number of new mature cells (RFP^+^-only) whereas expression a constitutively active Smox construct (Smox^SDVD^) increases the number relative to control. Expression of Smox^SDVD^ also expands the pool of stem and progenitor cells (GFP^+^, RFP^+^) relative to controls. Error bars represent SD; *p < 0.05, **p < 0.01, ***p < 0,001 and ****p < 0,0001

### Babo^A^ and Babo^C^ regulates different steps of ISC-to-EC differentiation

The *babo* gene encodes three different isoforms that differ in their extracellular domains and affinity for the three activin ligands. While BaboC is thought bind Daw, Babo^A^ was reported to mediate both Myo and Act-β signalling (Jensen et al. 2009; Awasaki et al. 2011; Song et al. 2017; Upadhyay et al. 2020). To analyse the role of each of these isoforms in regulating ISC-driven tissue turnover, we first analysed their expression levels. While isoforms A and C were expressed at intermediate and high levels, respectively, isoform B expression was hardly detectable in the gut (Fig. 2A). Knockdown of either Babo^A^ or BaboC reduced the number of ECs produced over time, showing that both isoforms are required for ISC proliferation and/or differentiation into ECs (Fig. 2B). By contrast, we did not observe any effect of Babo^B^ knockdown on tissue turnover rates (Fig. 2B). To analyse how BaboA and BaboC depletion affects ISC differentiation we made use of a cell fate sensor (Cfs^ts^) line allowing us to monitor the sizes of the ISC, EB and EC populations over time (Fig. 2C; (Martin et al. 2018)). Strikingly, knockdown of BaboA resulted in a depletion of EBs, while the ISC pool remained stable (Fig. 2D-E, G-G’’’), suggesting that BaboA is required in ISCs for their maturation into EBs. By contrast, knockdown of BaboC in ISCs/EBs caused an accumulation of both ISCs and EBs suggesting a defect in the maturation of EBs to ECs (Fig. 2D, F, G-G’’’). Indeed, EB-specific Babo^C^ depletion triggered an accumulation of EBs, showing that Babo^C^ is required in EBs to promote their maturation into ECs (Fig. 2H, J-K). As expected, EB-specific knockdown of BaboA had no effect on EBs numbers (Fig. 2H-I, K). Hence, Babo^A^ and Babo^C^ display different expression patterns and control different steps of ISC to EC differentiation.

**Figure 2:**
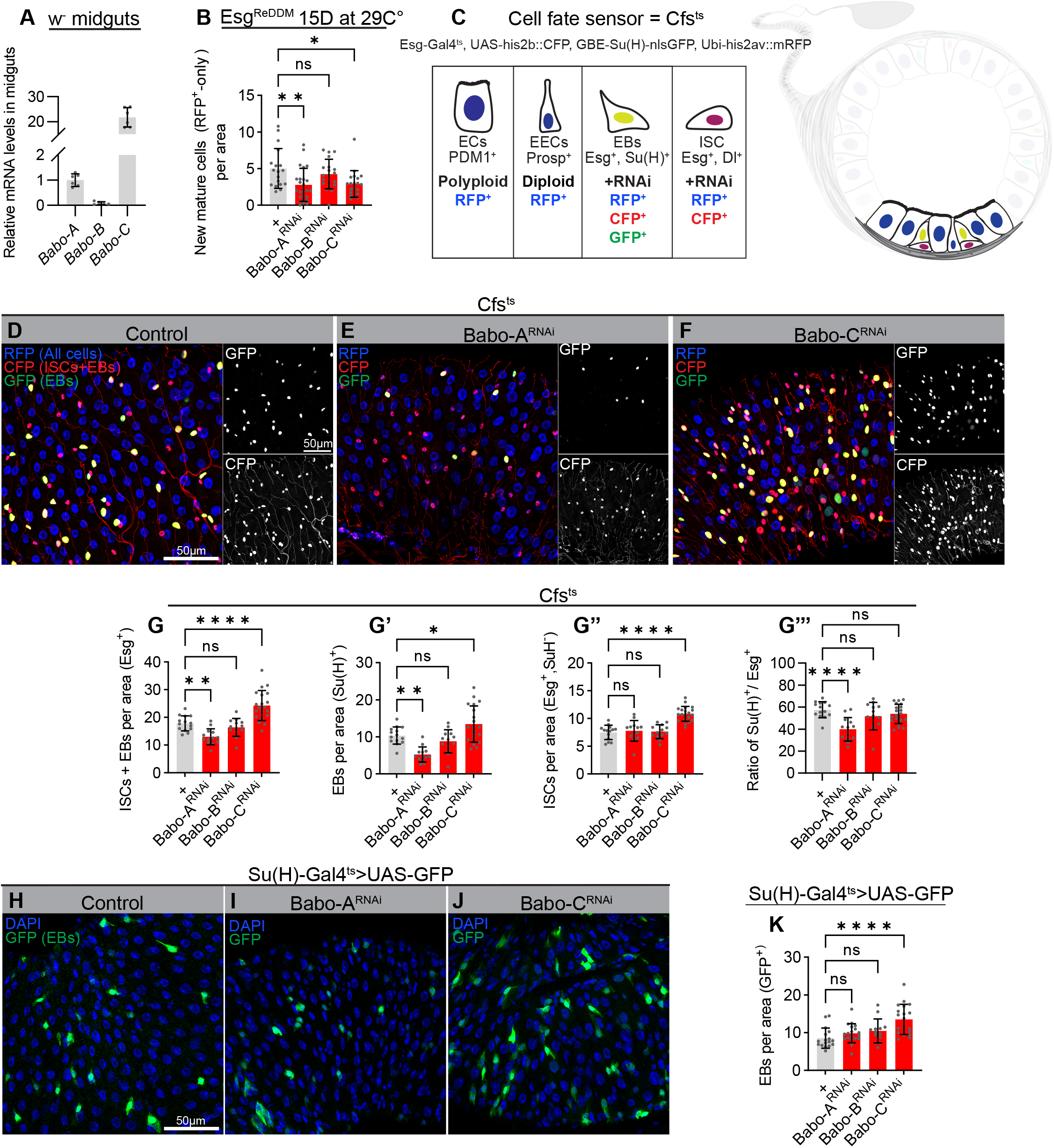
Babo isoforms A and C regulate different steps of ISC-to-EC differentiation. (A) RT-qPCR analysis on dissected control midguts showing differential expression of Babo-A, Babo-B and Babo-C isoforms. (B) Quantification of cell turnover in dissected posterior midguts from control flies and flies expressing Babo-A^RNAi^, Babo-B^RNAi^ and Babo-C^RNAi^ in stem and progenitor cells using Esg^ReDDM^ tracing for 15 days at homeostatic conditions. (C) Schematic diagram of the Cfs^ts^ system used to distinguish midgut cell types. Stem and progenitor cells are collectively visualized by expressing nuclear localized UAS-his2b::CFP with Escargot-Gal4, EBs are marked by nuclear GBE-Su(H)-nlsGFP while Ubi-his2av::mRFP causes all cells to express nuclear RFP. Thus, ISC and EBs are both marked by CFP, but only EBs are GFP-positive, whereas mature ECs and EECs are both marked by RFP but distinguished by nuclear size. (D-G’’’) Babo-A and Babo-C isoforms control distinct stages of ISC-EC differentiation. Representative confocal images of dissected posterior midguts from control flies (D) and flies expressing Babo-A^RNAi^ (E) or Babo-C^RNAi^ (F) in stem and progenitor cells using Cfs^ts^ and quantification of cell type numbers in each condition (G-G’’’). Depletion of Babo-A causes a loss of EBs, while Babo-C depletion increases the number of both ISCs and EBs relative to controls. (H-K) Babo-C isoform in EBs controls the EB-EC transition. Representative confocal images of dissected posterior midguts from control flies (H) and flies expressing Babo-A^RNAi^ (I) and Babo-C^RNAi^ (J) in EBs using Su(H)^ts^>UAS-GFP and quantification of EB numbers in each condition (K). Depletion of Babo-C causes an accumulation of EBs relative to controls. Error bars represent SD; *p < 0.05, **p < 0.01, ***p < 0,001 and ****p < 0,0001

### Act-β/Babo^A^ signaling promotes regenerative growth

The role of activin signalling in tissue repair has not been investigated in neither mammals nor flies. We therefore tested how tissue repair triggered by oral infection is affected by ISC/EB-specific Babo knockdown. Knockdown of all three Babo isoforms significantly reduced the infection-induced ISC proliferative response (Fig. 3A-B) and led to a decrease in organism viability after oral infection with pathogenic bacteria (Fig. 3C). The reduced regenerative response was recapitulated by ISC/EB-specific knockdown of Babo^A^, but not Babo^B^ or Babo^C^ (Fig. 3D). Using the cell fate sensor, we further showed that knockdown of BaboA in progenitor cells specifically reduces the number of EBs, whereas ISC numbers are slightly increased compared with infected control guts (Fig. 3E-G). Altogether, this is consistent with a key role of BaboA in promoting ISC divisions and ISC-to-EB differentiation. BaboA was previously reported to function as a receptor for both Act-β and Myo. However, while Act-β expression was strongly upregulated in response to oral infection (Fig. 3H), the expression of Myo was only moderately increased (Fig. S1A), suggesting that Act-β might be the principal driver of regenerative growth. We also observed an intermediate up-regulation of Daw 16 hours after infection (Fig. S1A’). In homeostatic conditions, Act-β expression is detected exclusively in EECs (Fig. 3I, K-K’; (Song et al. 2017)), but its expression is triggered in EBs in response to oral infection (Fig. 3I-J, K’). This is consistent with RNAseq data on FACS sorted gut cells showing a 100-fold upregulation of Act-β in EBs in response to oral PE infection (Dutta et al. 2015). In accordance with EBs being the main source of Act-β fuelling regenerative growth, we found that knockdown of Act-β in EBs, but not ISCs, EECs or ECs, significantly reduced the proliferative response (Fig. 3L-L’’’). Likewise, knockdown of BaboA diminished the progenitor pool in conditions of infection-induced regenerative growth and reduced tissue turnover rates (Fig. 3M-P). As BaboC promotes the maturation of EBs to ECs, knockdown of BaboC only had a modest effect on the progenitor pool (Fig. 3P’) but was required to promote EC replenishment in both homeostatic conditions and during tissue repair (Fig. 2K, Fig. 3M, O-P). In agreement with a role of activin signalling in promoting tissue repair, ISC/EB-specific knockdown of BaboA or EB-specific knockdown of Act-β reduced the resistance of flies to oxidative stress (Fig. S1B-B’).

**Figure 3:**
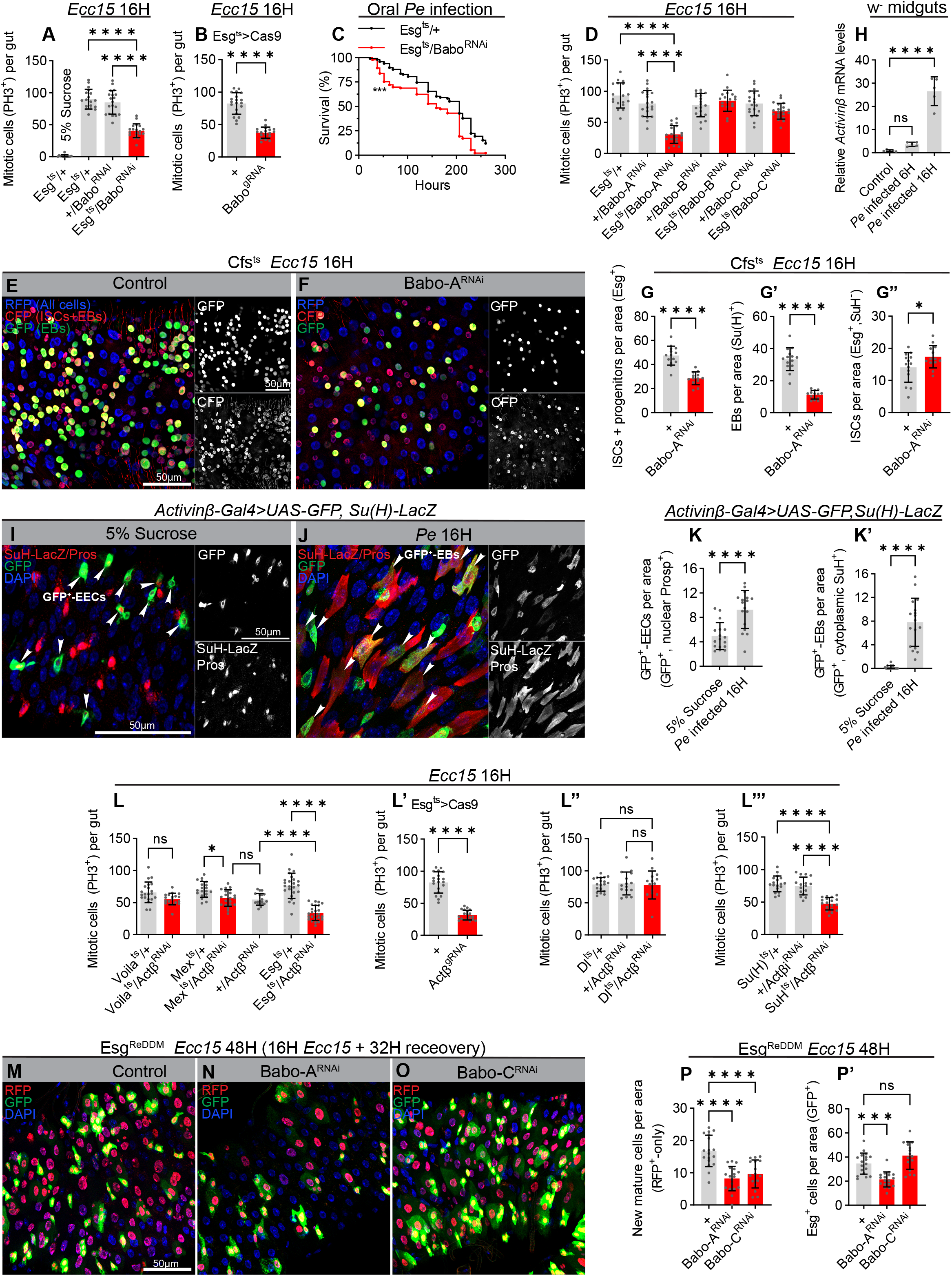
EB-derived Actβ supports regenerative growth and tissue repair. (A-B) Babo depletion in stem and progenitor cells reduces the proliferative the response. Quantification of mitotic cells (PH3^+^) in entire midguts in control flies and flies with ISC/EB-specific RNAi- (A) or CRISPR/Cas9-mediated (B) Babo knockdown 16 hours post oral infection with *Ecc15* or vehicle (5% sucrose). (C) Flies with ISC/EB-specific knockdown of Babo display reduced survival to oral infection. (D) ISC/EB-specific depletion of the Babo-A, but not Babo-B or Babo-C, isoform reduces the proliferative response to infection. Quantification of mitotic cells (PH3^+^) in entire midguts in control flies and flies expressing Babo-A^RNAi^, Babo-B^RNAi^ or Babo-C^RNAi^ in stem and progenitor cells using Esg^ts^ 16 hours post oral *Ecc15* infection. (E-G’’) Babo-A promotes the differentiation of ISCs to EBs. Representative confocal images of dissected posterior midguts from control flies (E) and flies expressing Babo-A^RNAi^ (F) in stem and progenitor cells using Cfs^ts^ 16 hours post oral *Ecc15* infection and quantification of cell type numbers. (H) RT-qPCR analysis on dissected control midguts 6- and 16 hours post oral *P.e* infection showing a strong upregulation of Act-β following infection. (I-K’) Actβ is upregulated in EBs during regeneration. Representative confocal images of dissected posterior midguts containing Actβ-Gal4>UAS-GFP and SuH-LacZ that marks EBs stained with β-Galactosidase (β-gal; cytoplasmic Red), Pros (nuclear Red), GFP (Green) and DNA (Blue) 16 hours post oral *Pe*. infection (J) or vehicle (5% sucrose) (I), and quantification of cell types positive for GFP-signal (K-K’). GFP is predominantly observed in EECs (nuclear Pros, white arrows) in unchallenged midguts (I), while *P.e* infection dramatically increases the number of GFP-positive EBs (cytoplasmic β-gal, white arrows (J). (L-L’’’) Knockdown of Actβ in EBs suppresses the proliferative response triggered by oral infection. Quantification of mitotic cells (PH3^+^) in entire midguts in control flies and flies expressing Actβ^RNAi^ in EECs (*voila^ts^*), ECs (*mex^ts^*), ISC+EBs (*esg^ts^*) (L) or Cas9+Actβ^gRNA^ in ISCs+EBs (L’) or flies expressing Actβ^RNAi^ in ISCs (Dl^ts^) (L’’) or EBs (Su(H)^ts^) (L’’’) 16 hours post oral *Ecc15* infection. (M-P’) Knockdown of Babo-A in ISCs/EBs slows down epithelial turnover. Representative confocal images of dissected posterior midguts from control flies (M) and flies expressing Babo-A^RNAi^ (N) and Babo-C^RNAi^ (O) in stem and progenitor cells 48 hours (16H on *Ecc15* + 32H recovery on normal food) post oral *Ecc15* infection using Esg^ReDDM^ tracing for 10 days and quantification of cell turnover in each condition (P-P’). Error bars represent SD; *p < 0.05, **p < 0.01, ***p < 0,001 and ****p < 0,0001

Many signals that stimulate proliferation and tissue repair in young animals also contribute to aged-related dysplasia. To investigate how augmented levels of activin signalling might affect longevity, we knocked down the activin inhibitor, FS, which was reported to counteract activin signalling in both flies and mammals. We observed an increase in ISC proliferation upon EC-specific knockdown of FS in both young and old animals accompanied by a reduced lifespan (Fig. S1C-D), suggesting that excess activin signalling might accelerate ageing-associated deterioration of the gut epithelium.

### Daw/Babo^C^ signaling adapts gut size to nutrient availability

The adult gut is a plastic organ that undergoes resizing in response to certain environmental cues. Hence, it was shown that the cycles of starvation and refeeding causes the gut to shrink and regrow, respectively, in a wide range of organisms (Altmann 1972; Andrew et al. 2015; Dunel-Erb et al. 2001; Lucchetta and Ohlstein 2017; McLeod et al. 2010; O’Brien et al. 2011; Secor, Stein, and Diamond 1994). Previous studies in flies have demonstrated an essential role of Dilp3 in driving progenitor and gut expansion following the first meals in newly eclosed flies (O’Brien et al. 2011). By contrast, it is not clear how short cycles of intermittent feeding affects progenitor number in the mature gut nor how adaptive growth is regulated in this condition. We therefore applied cycles of 48 hours of starvation followed by 24 hours of refeeding to 6 days old virgins. Two days of starvation was sufficient to shrink the gut, while 24 hours of refeeding allowed the gut to reach its pre-starvation size (Fig. 4A-B’). Interestingly, while 2 days of starvation decreased the total number of ECs, EECs, and ISCs, it increased EB numbers (Fig. 4C-C’’’). This contrasts with prolonged starvation, which results in a loss of EBs and massive apoptosis (McLeod et al. 2010; O’Brien et al. 2011). While we could detect moderate levels of apoptosis in ECs, we did not observe significant levels of apoptosis in EBs following 48 hours of starvation, suggesting that EBs are only lost upon prolonged starvation (Fig. 4D). The accumulation of EBs observed after 48 hours of starvation recapitulated the phenotype caused by EB-specific knockdown of BaboC (Fig. 2K), and hence, to test whether activin signalling regulates nutrient-dependent adaptive growth, we next monitored the expression levels of activin ligands during cycles of starvation and refeeding. Strikingly, we found that Daw expression was strongly suppressed by starvation and re-expressed upon refeeding (Fig. 4E). Although the expressions of both Act-β and Myo were slightly reduced upon nutrient deprivation, their expression levels were not restored upon refeeding (Fig. S1E-E’). Consistent with a role of Daw in coupling adaptive resizing of the gut with nutrient availability, knockdown of its receptor, BaboC, in progenitor cells prevented normalisation of EB numbers (Fig. 4F-K) and impaired resizing of the gut in refed animals (Fig. 4J-J’’). Hence, starvation-mediated repression of Daw signalling might shrink the gut by preventing EB differentiation, while refeeding promotes Daw-dependent EB maturation and adaptive growth. Consistent with this, Daw is required in ECs, where it is also highly expressed (Fig. 4L), to resize the gut following refeeding by permitting the accumulated pool of EBs to mature into ECs (Fig. 4M-M’). Intermittent cycles of starvation might be a frequently occurring phenomenon in the nature, and hence, loss of gut plasticity might have far-reaching consequences for survival. To test this, we subjected flies to repetitive cycles of fasting and refeeding and monitored survival. Indeed, knockdown of BaboC in progenitor cells is sufficient to make flies more sensitive to successive cycles of starvation/refeeding (Fig. 4N), demonstrating the physiological importance of Daw/Babo^c^ signalling in coupling nutrient intake with adaptive gut resizing.

**Figure 4:**
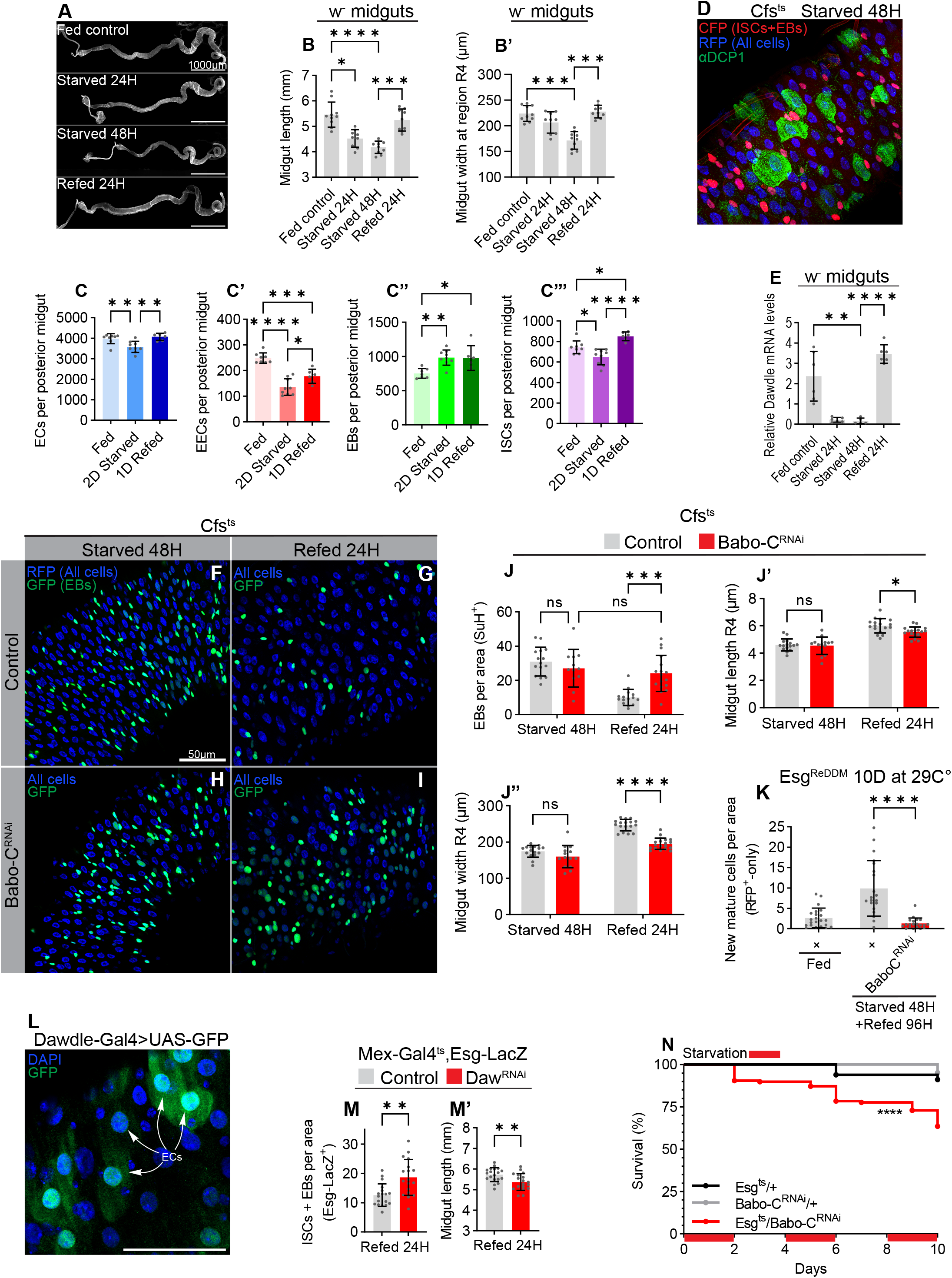
Daw couples calorie intake with EB differentiation to promote adaptive growth. (A-B’) Representative images of dissected midguts of fed, 24H starved, 48H starved and 24H refed controls and quantification of midgut length (B) and width (B’) in each conditon. Starvation results in a gradual decrease in gut size that is completely reversible by refeeding for 24 hours. (C-C’’’) Starvation triggers an accumulation of EBs. Quantification of the total number of cells in the posterior midgut of fed, 48H starved and 48H starved + 24H refed animals using Cfs^ts^. (D) Representative image of dissected midguts of 48H starved Cfs^ts^ animals using Cfs^ts^ stained for active caspases (Dcp1, in green) to visualize apoptotic cells. Apoptosis is observed in ECs (large cells) and EECs (small CFP^-^ cells) but not in EBs. (E) RT-qPCR analysis on dissected control midguts in fed, 24H starved, 48H starved and 48 H starved + 24H refed flies. Expression of *Daw* dynamically responds to nutrient availability. (F-I) Babo-C is required for adaptive growth. Representative confocal images of dissected posterior midguts from control flies (F-G) or flies with ISC/EB-specific knockdown of Babo-C^RNAi^ using Cfs^ts^ (H-I) starved for 48 hours (F, H) and refed for 24 hours (G, I), and quantification of cell type numbers (J) and midgut dimensions (J-J’’) in each condtion. (K) Quantifcation of cell turnover in dissected posterior midguts from fed, 48H starved, or 48H starved + 48H refed control flies or flies with ISC/EB-specific knockdown of Babo-C^RNAi^ using Esg^ReDDM^ tracing for 10 days. (L) Daw is expressed in ECs. Representative confocal image of dissected prosterior midgut from an adult fly bearing Daw-Gal4>UAS-nlsGFP stained for GFP (Green) and DNA (Blue) at homeostatic conditions. GFP expression is observed in large polyploidy ECs. (M-M’) EC-derived Daw supports EB differentiation during adaptive growth. Quantification of midgut lengths and ISC + EB numbers (using *esg*-LacZ) in dissected posterior midguts from 48H starved + 24H refed control flies or flies with EC-specific Daw knockdown. Gut resizing and normalization of ISC + EB numbers are decreased upon Daw depletion in ECs. (N) Flies with ISC/EB-specific knockdown of BaboC show reduced resistance to successive cycles of starvation/refeeding. Error bars represent SD; *p < 0.05, **p < 0.01, ***p < 0,001 and ****p < 0,0001

## Discussion

In mammals, the BMP branch of the TGF-β superfamily has a well-defined role in regulating intestinal homeostasis. BMP signaling exhibits a gradient along the crypt to villus axis. Low levels of BMP signaling at the bottom of the crypt is essential to promote Lgr5^+^ ISC selfrenewal, while high levels of BMP suppress proliferation and prevents expansion of the Lgr5^+^ ISC population outside of the crypt (Gehart and Clevers 2019; Wang and Chen 2018). The role of BMP/Dpp signaling in the fly gut is more complex, as different sources of Dpp was reported to promote (Ayyaz, Li, and Jasper 2015; Tian, Wang, and Jiang 2017) or restrict (Guo, Driver, and Ohlstein 2013; Zhou et al. 2015) regenerative growth. Interestingly, Aayez et al reported that Smox, normally considered an activin-specific mediator, was required downstream of hemocyte-derived Dpp to trigger regenerative growth (Ayyaz, Li, and Jasper 2015). This was contradicted by a more recent study showing that hemocytes are dispensable for proliferative response triggered by intestinal infection, bringing into question whether Dpp mediates Smox-dependent regenerative growth (Chakrabarti et al. 2016). According to the data presented here, an alternative possibility is that Smox-dependent regenerative growth is triggered by activins.

The intense research put into deciphering the role of BMP signaling in maintaining gut homeostasis contrasts with a lack of studies addressing the function of nodal/activin signaling in the adult gut. Here we show that activin signaling plays a key role in controlling epithelial turnover rates in the gut by controlling ISC proliferation and fate choices. While Act-β signals through BaboA to promote ISC self-renewal and their differentiation into EBs, Daw/Babo^C^ signaling is required in EBs for their maturation into ECs. Notably, Daw and Act-β expression levels are regulated by different environmental cues. Intestinal infections induce high levels of Act-β expression, while Daw expression is only moderately affected. This is consistent with the observation that EB-derived Act-β is required for the accelerated epithelial turnover rates associated with regenerative growth. The observation that Daw expression is only moderately changed by intestinal infections, suggest that basal levels of Daw suffice to promote EB maturation in this condition. Interestingly, Act-β produced by the PNS acts on the hematopoietic progenitors in hematopoietic pockets to stimulate their proliferation in larvae (Makhijani et al. 2017). As the prime function of the PNS is to detect innocuous and noxious sensory stimuli, it is possible that activin signalling has a more general role in coupling harmful environmental cues with progenitor expansion to promote host defence. As mentioned earlier, Daw is related to vertebrate TGFβ1. In mammals, TGF-β/TβR1 signaling gradually increases from the crypt towards the tip of the villi with nuclear Smad3 being detectable in late TA and differentiated cells (Fink and Wrana 2022). Based on *in vitro* and *in vivo* experiments, TGF-β is thought to promote terminal differentiation of ECs as these move out the crypt and up the villi (Cammareri et al. 2017; Flentjar et al. 2007; Halttunen et al. 1996; Kurokowa, Lynch, and Podolsky 1987; Yamada et al. 2013). Moreover, TGF-β signaling is required for tissue repair, as epithelial wide inactivation of TβRII results in the accumulation of undifferentiated progenitor cells (Oshima et al. 2015). Our data suggest that the role of Daw/TGFβ signaling in promoting terminal EC differentiation might be conserved between flies and mammals.

The rapid tissue turnover observed in mammalian guts is an energy expensive process, consuming up to 15% of the total basal metabolic rate (Aiello and Wheeler 1995), and hence, during periods with reduced food intake, it is likely advantageous to reduce the total number of cells needing constant replenishment. Indeed, in both flies and mammals, prolonged periods of starvation can result in a reduction in gut size of up to 50%, which can be recovered following a few days of refeeding. In mammals, starvation slows down the proliferation in the transient amplifying region thereby reducing the number of epithelial cells, while +4 ISCs numbers and niche function is preserved (Richmond et al. 2015; Yilmaz et al. 2012). This allows for rapid expansion of the epithelium upon refeeding. In flies, the expansion of progenitor cells in newly eclosed flies is triggered by food intake and mediated by the insulinlike molecule Dilp3 (O’Brien et al. 2011). By contrast, it is not clear how adaptive growth is triggered during cycles of starvation-refeeding in mature guts. So far, most starvation-refeeding studies have been performed within 2 days of eclosion, when progenitor cells are still sparse (Lucchetta and Ohlstein 2017; O’Brien et al. 2011; Bonfini et al. 2021), or involves prolonged periods of protein starvation of 7-15 days, which triggers high levels of apoptosis and dramatically reduces total numbers of EBs and ECs (McLeod et al. 2010). Importantly, as a proxy for intermittent feeding opportunities encountered by flies in the nature, it might be physiologically more relevant to study how shorter periods of starvation affects progenitor cell numbers. Here, we show that 24-48 hours of starvation, a condition that is not associated with massive cell death, results in an accumulation of EBs, despite a reduction in ECs and total number of cells. While the failure of EBs to differentiate into ECs results in shrinkage of the gut, it also generates a pool of poised EBs committed to promote adaptive growth upon refeeding. Indeed, EB numbers normalize shortly after refeeding, a process that depends on Daw/Babo^C^ signaling. Accordingly, Daw expression is strongly repressed upon nutrient deprivation, which likely restricts EB to EC differentiation, and reinduced upon refeeding. All together this suggests that Daw/BaboC signaling is essential for coupling adaptive resizing of the gut with nutrient intake. In mammals, nutrient restriction results in an accumulation of dormant ISCs (d-ISCs), also called +4 ISCs as they mark the transition from rapidly cycling Lgr5+ ISCs at the base of the crypt and the TA zone (Richmond et al. 2015). While d-ISCs cycle slowly or not at all in homeostatic conditions, they accumulate in response to nutrient restriction and participate in the regenerative growth during refeeding (Richmond et al. 2015). Hence, rapid regrowth of the gut upon calorie intake might rely on differentiation of stalled progenitors in both flies and mammals. The Daw-mediated adaptive resizing of the gut is physiologically relevant, as knockdown of Babo^C^ in EBs decreases the chances of survival of flies exposed to intermittent cycles of starvation and refeeding. Future research should aim at addressing whether TGF-β signaling plays a similar prominent role in adaptation of gut size to calorie intake in mammals.

While regulators of organ plasticity are essential for host adaptation, reproductivity and survival in an ever-changing environment, the same signals are often deregulated in cancers. Indeed, mutations affecting TGF-β/Nodal/activin signaling has been observed in a wide range of cancers, and better understanding of how TGF-β/Nodal/activin signaling affects intestinal growth might provide a starting point for the development of therapeutic strategies targeting colorectal cancers.

## Acknowledgement

We are grateful to all members of the Colombani and Andersen laboratory for scientific discussion and for carefully reading the manuscript. We thank L. O’Brien, M. Dominguez, M. O’Connor, A. Gallet, the Bloomington Stock Center and the Vienna Drosophila RNAi Center for fly stocks. J.C. and D.S.A. are funded by H2020 European Research Council grant number 803630, Novo Nordisk Foundation grant number NNF180C0033920. We thank the Carlsberg foundation for an equipment grant CF19-0353.

## Author contribution

C.F.C., J.C. and D.S.A designed the research, C.F.C., Q.L, R.L, J.C., and D.S.A performed the genetic screen, C.F.C. conducted all the remaining experiments for the manuscript, C.F.C., J.C. and D.S.A analyzed the data, and J.C. and D.S.A supervised the project, and D.S.A. wrote the manuscript.

---

**Supplementary Figure 1:**
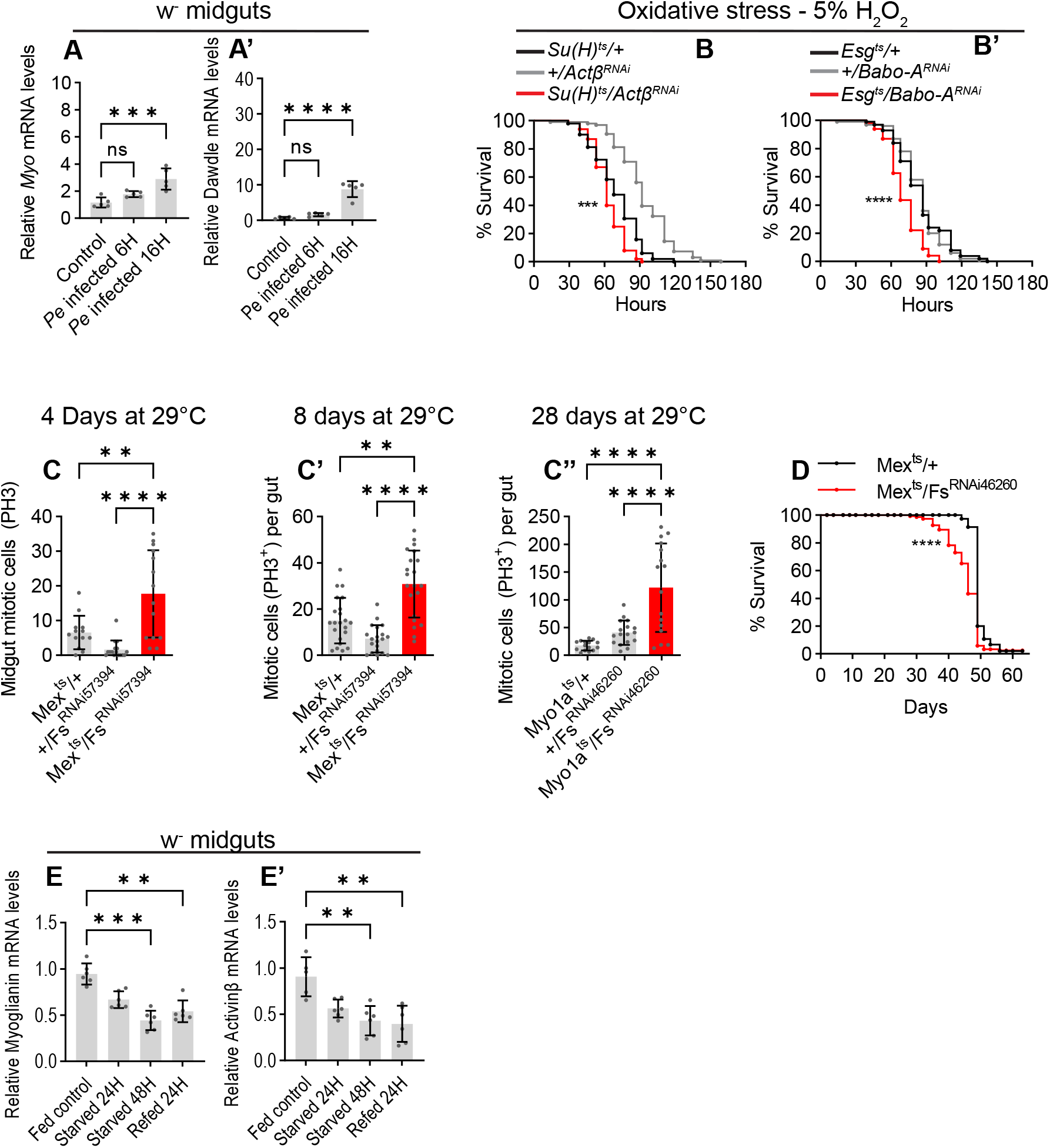
(A-A’) RT-qPCR analysis on dissected control midguts 6- and 16 hours post oral *P.e* infection showing upregulation of *Myo* and *Daw* following infection. (B-B’) Flies with depletion of Act-β in EBs using SuH^ts^ or Babo-A in stem and progenitor cells using Esg^ts^ display reduced survival to oxidative stress triggered by ingestion of H_2_O_2_. (C-C’’) Quantification of mitotic cells (PH3^+^) in entire midguts in control flies and flies expressing Fs^RNAi^ (BL57394 or v46260) in ECs using Mex^ts^ or Myo^ts^ after 4 (C), 8 (C’), or 28 (C’’) days of RNAi induction. (D) EC-specific depletion of Fs (Red) shortens lifespan relative to control flies (Black). (E-E’) RT-qPCR analysis on midguts dissected from fed, 24H starved, or 48H starved + 24H refed Flies. Error bars represent SD; *p < 0.05, **p < 0.01, ***p < 0,001 and ****p < 0,0001

## Materials and methods

### Fly stocks and husbandry

Animals were maintained on a standard cornmeal diet (containing: 82 g/L cornmeal, 60 g/L sucrose, 34g/L yeast, 8 g/L agar, 4.8 mL/L propionic acid and 1.6 g/L methyl-4-hydroxybenzoate) at 25 °C and 60% relative humidity under 12-hour light/dark cycle conditions. Virgin females were used for all experiments unless otherwise stated. w^1118^ was used as control. Temporal control of transgene expression using tubGal80ts in adult flies was achieved by raising flies at 18°C through development until 4-7 days after eclosion to allow maturation of the digestive system. Then, flies were shifted to 29°C to induce Gal4 mediated transgene expression in a cell-type specific manner. Duration of UAS induction was 8-10 days unless otherwise stated. Flies were flipped onto fresh medium every second day. For experiments using CRISPR/Cas9, UAS-Cas9.P2 and UAS-gRNA was co-induced for 10 days to allow mutagenesis of the target gene. Then, flies were shifted to 18°C for an additional 30 days to eliminate the Cas9 protein and re-establish a homeostatic state with the targeted gene removed.

The following lines were generous gifts from the colleagues in the fly community: cell fate sensor, cfs: Esg-Gal4, UAS-CFP,Su(H)Gbe-GFP;ubi-his2av-RFP (Lucy O’Brien, Stanford University, USA). Myo1a-Gal4, Su(H)Gbe-Gal4 and Voila-Gal4 (Armel Gallet, ISA, France). Esg ReDDM: Esg-Gal4, UAS-CD8::GFP; UAS-H2B::RFP, tubGal80ts/TM6b (Maria Dominguez, IN, Spain). UAS-dSmad2^SDVD^, Actβ-Gal4 and Daw-Gal4 (Michael O’Connor, CBS, USA). Actβ-gRNA v342095, Babo-gRNA v341225, dSmad2-gRNA v342177 and UAS-Fs-RNAi v46260 were obtained from the Vienna Drosophila RNAi center. UAS-Actβ-RNAi BL29597, UAS-Daw-RNAi BL34974, UAS-Babo-RNAi BL29533, UAS-BaboA-RNAi BL44400, UAS-BaboB-RNAi BL44401, UAS-BaboC-RNAi BL44402, UAS-dSmad2-RNAi BL41670, UAS-Fs-RNAi BL57394, Babo-Gal4 BL83164, Esg-Gal4 BL93857, Su(H)Gbe-LacZ BL83352, esg-lacZ BL10359, UAS-Cas9.P2 BL58986, UAS-GFP BL39760, tub-Gal80ts BL7107 and BL7108, Mex-Gal4 BL91368 were obtained from the Bloomington stock center.

### Oral infections

For oral infections with bacteria (Ecc15 or Pe), overnight cultures were grown from single colonies (Ecc15) or directly from a glycerol stock kept at −80°C (Pe). Cultures were grown in conical flasks containing LB broth at 29°C (Ecc15) or at 30°C with rifampicin supplied (Pe). The next day optical density OD_600_ was measured, cultures were spun down and the remaining bacterial pellet was resuspended in an appropriate volume of 10% sucrose such that OD_600_ was adjusted to 200 in 5% sucrose. The concentrated bacterial solution was then, in a volume of 50 μl, added directly onto Whatman filter paper discs resting on the surface of normal fly food with reduced yeast content (containing: 82 g/L cornmeal, 60 g/L sucrose, 17g/L yeast, 8 g/L agar and 4.6 g/L methyl-4-hydroxybenzoate). To ensure efficient intake of bacteria, flies were sorted 10 by 10 in empty vials and starved for 2-3 hours before transfer to the prepared vials containing bacteria.

### Starvation refeeding scheme

Flies were nutrient starved in vials containing H2O + 1% agar for 24 or 48 hours. For refeeding, 48 hours starved flies were transferred to normal food and allowed to feed for 24 hours.

### Longevity

Flies raised at 18°C through development were allowed to emerge over 48 hours and transferred to fresh media for an additional 48 hours at 25°C to mate. Mated female flies were then sorted into vials containing sugar/yeast food (consisting of 90 g/L sucrose, 80 g/L yeast, 10 g/L agar, 0,5% propionic acid and 0,15% methyl-4-hydroxybenzoate) and transferred to 29°C to induce transgenes. Flies were transferred onto fresh food every 2-3 day and the number of dead animals was assessed each 2-3 days until all animals had died. UAS-RNAi transgenes were backcrossed for at least 7 generations into the w^1118^ background used as control.

### Oxidative stress survival

To assess survival to oxidative stress, adult virgin female flies were kept at 18°C for 4 days before being transferred to 29°C for 15 days. Flies were then sorted 10 by 10 and starved for 2-3 hours at 29°C in empty vials prior to oxidative stress challenge. Sugar/yeast food supplemented with 5% H_2_O_2_ was freshly prepared on the day. Following starvation, flies were transferred to H_2_O_2_ food and kept on the same media and the number of dead animals was assessed three times a day until all animals had died.

### Dissections and immunohistochemistry

Midguts were dissected in PBS and immediately transferred into fixative consisting of 4% paraformaldehyde in PBS for 1 hour at RT. Fixed midguts were washed twice with PBS and further two times 15 min with PBS + 0,1% Trition (PBS-T) with agitation. Midguts that were not stained with antibodies (EsgReDDM and Cfs^ts^) were immediately washed once with PBS and mounted on microscopy slides at this stage. Midguts that were stained with antibodies were incubated for 2 hours in blocking solution (PBS-T containing 10% FCS) followed by incubation with primary antibodies prepared in blocking solution at 4°C overnight. The next day, midguts were washed three times 15 min with PBS-T and incubated with secondary antibodies for 3 hours at RT or overnight at 4°C. At last, the stained midguts were washed two times 15 min with PBS-T followed by a single wash with PBS before being mounted on glass slides in Vectashield mounting media with or without DAPI. All midguts were mounted with 0,12mm SecureSeal spacers (Grace Bio-Labs) except for experiments assessing the number of mitotic (PH3+) cells per gut. Primary antibodies were used with the following dilutions: rabbit anti-PH3 1:1000 (Milipore 06-570), chicken anti-GFP 1:10000 (Abcam 13970), mouse anti-β-Galactosidase (DSHB 40-1A), mouse anti-Armadillo 1:100 (DSHB N2-7A1), mouse anti-prospero 1:200 (DSHB MR1A) and mouse anti-Delta 1:100 (DSHB C594.9B). Secondary antibodies were used with the following dilutions: Alexa Fluor 488-conjugated goat antirabbit 1:1000 (Thermofisher #A-11008), Alexa fluor 488-conjugated goat anti-chicken 1:1000 (Thermofisher #A-21467) and Cy3-conjugated goat anti-mouse 1:1000 (Thermofisher #A-10521)

### Microscopy

Mounted midguts were imaged on an inverted Zeizz LSM-900 confocal microscope with Zen Blue software using 5x, 20x, or 40x objectives. For imaging CFP, GFP and RFP together (Cfsts), the following bandwidth settings were used to avoid spectral bleed-through: CFP excitation at 405 nm and detection at 400-480 nm, GFP excitation at 488 nm and detection at 510-550 nm, RFP excitation at 561 nm and detection at 580-700 nm. 6-12 confocal stacks were acquired with an interval of 2 μm spanning the intestinal epithelium from the base to the lumen. An identical number of z-stacks were acquired for each experiment involving quantitative assessment of cell types. All images including the ones for quantifications were acquired in the posterior gut in region R4bc whereas the image in fig. 1A is taken in region R5. Images were processed using Fiji and Adobe Photoshop software.

### Image analysis and quantifications

Image analysis and quantifications were performed using the open-source FIJI software. For quantitative measurements of specific cell types per area, a z-stacks were acquired with a 20x objective in the R4bc region for each mounted gut. Acquired z-stacks were converted to maximum-intensity projections and cell numbers was manually counted using the Cell Counter Fiji plugin and normalized to the total epithelial area counted in. Cells per area is defined as cells per 10000 μm2 in all cases. For quantifications of gut dimensions, tiled images of entire fixed midguts mounted with 0,12mm spacers were acquired with fluorescence imaging with a 5x objective and stitched together in Zen Blue imaging software (Zeizz). Length was then measured in Fiji by drawing a segmented line using spline along the center of each midgut while R4 width was measured by drawing a straight line perpendicular to the midgut at the widest point in the R4 region.

### RNA extraction and qPCR

8-10 dissected midguts per biological replicate was directly transferred into lysis buffer and flash frozen in liquid nitrogen. Total RNA was extracted using RNAeasy microkit (Qiagen) according to the manufacturer’s instructions. For cDNA synthesis, RNA was treated with DNase and reverse-transcribed using Superscript II reverse transcriptase (Invitrogen). The resulting cDNA was used for real-time RT-PCR on a QuantStudio 5 Real-Time PCR system using RealQ Plus 2x Master Mix Green (Ampliqon) with 8 ng of cDNA template. Samples were normalized to levels of ribosomal protein 49 expression levels and analyzed with QuantStudio 5 Real-Time PCR software using the delta-delta Ct method. Five to six biological replicates were used for each condition or genotype and triplicate measurements were performed. The following primers were used:

Actβ _F 5’-ACG GCA AAT TTT GAC AAA GC-3’
Actβ_R 5’-TTG GTA TCA TTC GTC CAC CA-3’
Daw _F 5’-TGA GCC ACC TCA TCC AAA TCA CCT-3’
Daw_R 5’-TCG ATC ACG ATG AAT GGC CGG TAA-3’
Myo _F 5’-GGC GAC CAC ATA ATG ACT-3’
Myo_R 5’-TTA GCA TCA TCT CCC TGC ATT-3’
Babo-A_F 5’-GGC TTT TGT TTC ACG TCC GTG GA-3’
Babo-A_R 5’-CTG TTT CAA ATA TCC TCT TCA TAA TCT CAT-3’
Babo-B_F 5’-GCA AGG ACA GGG ACT TCT G-3’
Babo-B_R 5’-GGC ACA TAA TCT TGG ACG GAG-3’
Babo-C_F 5’-GAC CAG TTG CCA CCT GAA GA-3’
Babo-C_R 5’-TGG CAC ATA ATC TGG TAG GAC A-3’

### Statistics

Visualization of data in graphs and statistical analysis were performed using GraphPad Prism software. Each dataset was assessed for normal distribution using Shapiro-Wilks normality test before analysis. Comparisons of 3 or more groups were analyzed by one-way ANOVA followed by Tukey’s multiple comparison tests if passing normality tests. If similar comparisons did not pass normality tests, Kruskal-Wallis test followed by Dunn’s multiple comparison tests were applied. Differences between control and one group was analyzed by Student’s t-test if passing normality tests or Mann-Whitney test if not. All error bars indicate standard deviation. Comparison of survival curves was analyzed using Mantel-Cox Log Rant test. p values are indicated as follows: *p < .05, **p < 0.01, ***p < 0,001 and ****p < 0,0001.

## References

Aiello, L.C., and P. Wheeler. 1995. ‘The expensive-tissue hypothesis’, Curr. Anthropol., 36: 199–221.

Altmann, G. G. 1972. ‘Influence of starvation and refeeding on mucosal size and epithelial renewal in the rat small intestine’, Am J Anat, 133: 391–400.

Andrew, A. L., D. C. Card, R. P. Ruggiero, D. R. Schield, R. H. Adams, D. D. Pollock, S. M. Secor, and T. A. Castoe. 2015. ‘Rapid changes in gene expression direct rapid shifts in intestinal form and function in the Burmese python after feeding’, Physiol Genomics, 47: 147–57.

Antonello, Z. A., T. Reiff, E. Ballesta-Illan, and M. Dominguez. 2015. ‘Robust intestinal homeostasis relies on cellular plasticity in enteroblasts mediated by miR-8-Escargot switch’, EMBO J, 34: 2025–41.

Awasaki, T., Y. Huang, M. B. O’Connor, and T. Lee. 2011. ‘Glia instruct developmental neuronal remodeling through TGF-beta signaling’, Nat Neurosci, 14: 821–3.

Ayyaz, A., H. Li, and H. Jasper. 2015. ‘Haemocytes control stem cell activity in the Drosophila intestine’, Nat Cell Biol, 17: 736–48.

Biteau, B., and H. Jasper. 2014. ‘Slit/Robo signaling regulates cell fate decisions in the intestinal stem cell lineage of Drosophila’, Cell Rep, 7: 1867–75.

Bonfini, A., A. J. Dobson, D. Duneau, J. Revah, X. Liu, P. Houtz, and N. Buchon. 2021. ‘Multiscale analysis reveals that diet-dependent midgut plasticity emerges from alterations in both stem cell niche coupling and enterocyte size’, Elife, 10.

Brummel, T., S. Abdollah, T. E. Haerry, M. J. Shimell, J. Merriam, L. Raftery, J. L. Wrana, and M. B. O’Connor. 1999. ‘The Drosophila activin receptor baboon signals through dSmad2 and controls cell proliferation but not patterning during larval development’, Genes Dev, 13: 98–111.

Cammareri, P., D. F. Vincent, M. C. Hodder, R. A. Ridgway, C. Murgia, M. Nobis, A. D. Campbell, J. Varga, D. J. Huels, C. Subramani, K. L. H. Prescott, C. Nixon, A. Hedley, S. T. Barry, F. R. Greten, G. J. Inman, and O. J. Sansom. 2017. ‘TGFbeta pathway limits dedifferentiation following WNT and MAPK pathway activation to suppress intestinal tumourigenesis’, Cell Death Differ, 24: 1681–93.

Chakrabarti, S., J. P. Dudzic, X. Li, E. J. Collas, J. P. Boquete, and B. Lemaitre. 2016. ‘Remote Control of Intestinal Stem Cell Activity by Haemocytes in Drosophila’, PLoS Genet, 12: e1006089.

Colombani, J., and D. S. Andersen. 2020. ‘The Drosophila gut: A gatekeeper and coordinator of organism fitness and physiology’, Wiley Interdiscip Rev Dev Biol, 9: e378.

de Navascues, J., C. N. Perdigoto, Y. Bian, M. H. Schneider, A. J. Bardin, A. Martinez-Arias, and B. D. Simons. 2012. ‘Drosophila midgut homeostasis involves neutral competition between symmetrically dividing intestinal stem cells’, EMBO J, 31: 2473–85.

Dunel-Erb, S., C. Chevalier, P. Laurent, A. Bach, F. Decrock, and Y. Le Maho. 2001. ‘Restoration of the jejunal mucosa in rats refed after prolonged fasting’, Comp Biochem Physiol A Mol Integr Physiol, 129: 933–47.

Dutta, D., N. Buchon, J. Xiang, and B. A. Edgar. 2015. ‘Regional Cell Specific RNA Expression Profiling of FACS Isolated Drosophila Intestinal Cell Populations’, Curr Protoc Stem Cell Biol, 34: 2F 2 1–2F 2 14.

Ellis, J. E., L. Parker, J. Cho, and K. Arora. 2010. ‘Activin signaling functions upstream of Gbb to regulate synaptic growth at the Drosophila neuromuscular junction’, Dev Biol, 342: 121–33.

Fink, M., and J. L. Wrana. 2022. ‘Regulation of homeostasis and regeneration in the adult intestinal epithelium by the TGF-beta superfamily’, Dev Dyn.

Flentjar, N., P. Y. Chu, A. Y. Ng, C. N. Johnstone, J. K. Heath, M. Ernst, P. J. Hertzog, and M. A. Pritchard. 2007. ‘TGF-betaRII rescues development of small intestinal epithelial cells in Elf3-deficient mice’, Gastroenterology, 132: 1410–9.

Gehart, H., and H. Clevers. 2019. ‘Tales from the crypt: new insights into intestinal stem cells’, Nat Rev Gastroenterol Hepatol, 16: 19–34.

Guo, Z., I. Driver, and B. Ohlstein. 2013. ‘Injury-induced BMP signaling negatively regulates Drosophila midgut homeostasis’, J Cell Biol, 201: 945–61.

Guo, Z., and B. Ohlstein. 2015. ‘Stem cell regulation. Bidirectional Notch signaling regulates Drosophila intestinal stem cell multipotency’, Science, 350.

Halttunen, T., A. Marttinen, I. Rantala, H. Kainulainen, and M. Maki. 1996. ‘Fibroblasts and transforming growth factor beta induce organization and differentiation of T84 human epithelial cells’, Gastroenterology, 111: 1252–62.

Herrera, S. C., D. Sainz de la Maza, L. Grmai, S. Margolis, R. Plessel, M. Burel, M. O’Connor, M. Amoyel, and E. A. Bach. 2021. ‘Proliferative stem cells maintain quiescence of their niche by secreting the Activin inhibitor Follistatin’, Dev Cell, 56: 2284894 e6.

Hu, D. J., and H. Jasper. 2019. ‘Control of Intestinal Cell Fate by Dynamic Mitotic Spindle Repositioning Influences Epithelial Homeostasis and Longevity’, Cell Rep, 28: 2807–23 e5.

Jensen, P. A., X. Zheng, T. Lee, and M. B. O’Connor. 2009. ‘The Drosophila Activin-like ligand Dawdle signals preferentially through one isoform of the Type-I receptor Baboon’, Mech Dev, 126: 950–7.

Kadaja, M., B. E. Keyes, M. Lin, H. A. Pasolli, M. Genander, L. Polak, N. Stokes, D. Zheng, and E. Fuchs. 2014. ‘SOX9: a stem cell transcriptional regulator of secreted niche signaling factors’, Genes Dev, 28: 328–41.

Kurokowa, M., K. Lynch, and D. K. Podolsky. 1987. ‘Effects of growth factors on an intestinal epithelial cell line: transforming growth factor beta inhibits proliferation and stimulates differentiation’, Biochem Biophys Res Commun, 142: 775–82.

Lengil, T., D. Gancz, and L. Gilboa. 2015. ‘Activin signaling balances proliferation and differentiation of ovarian niche precursors and enables adjustment of niche numbers’, Development, 142: 883–92.

Liu, X., J. J. Hodgson, and N. Buchon. 2017. ‘Drosophila as a model for homeostatic, antibacterial, and antiviral mechanisms in the gut’, PLoS Pathog, 13: e1006277.

Lonardo, E., P. C. Hermann, M. T. Mueller, S. Huber, A. Balic, I. Miranda-Lorenzo, S. Zagorac, S. Alcala, I. Rodriguez-Arabaolaza, J. C. Ramirez, R. Torres-Ruiz, E. Garcia, M. Hidalgo, D. A. Cebrian, R. Heuchel, M. Lohr, F. Berger, P. Bartenstein, A. Aicher, and C. Heeschen. 2011. ‘Nodal/Activin signaling drives self-renewal and tumorigenicity of pancreatic cancer stem cells and provides a target for combined drug therapy’, Cell Stem Cell, 9: 433–46.

Lucchetta, E. M., and B. Ohlstein. 2017. ‘Amitosis of Polyploid Cells Regenerates Functional Stem Cells in the Drosophila Intestine’, Cell Stem Cell, 20: 609–20 e6.

Makhijani, K., B. Alexander, D. Rao, S. Petraki, L. Herboso, K. Kukar, I. Batool, S. Wachner, K. S. Gold, C. Wong, M. B. O’Connor, and K. Bruckner. 2017. ‘Regulation of Drosophila hematopoietic sites by Activin-beta from active sensory neurons’, Nat Commun, 8: 15990.

Martin, J. L., E. N. Sanders, P. Moreno-Roman, L. A. Jaramillo Koyama, S. Balachandra, X. Du, and L. E. O’Brien. 2018. ‘Long-term live imaging of the Drosophila adult midgut reveals real-time dynamics of division, differentiation and loss’, Elife, 7.

Massague, J. 2008. ‘TGFbeta in Cancer’, Cell, 134: 215–30.

McLeod, C. J., L. Wang, C. Wong, and D. L. Jones. 2010. ‘Stem cell dynamics in response to nutrient availability’, Curr Biol, 20: 2100–5.

Micchelli, C. A., and N. Perrimon. 2006. ‘Evidence that stem cells reside in the adult Drosophila midgut epithelium’, Nature, 439: 475–9.

Min, S., A. Oyelakin, C. Gluck, J. E. Bard, E. C. Song, K. Smalley, M. Che, E. Flores, S. Sinha, and R. A. Romano. 2020. ‘p63 and Its Target Follistatin Maintain Salivary Gland Stem/Progenitor Cell Function through TGF-beta/Activin Signaling’, iScience, 23: 101524.

Ng, J. 2008. ‘TGF-beta signals regulate axonal development through distinct Smad-independent mechanisms’, Development, 135: 4025–35.

O’Brien, L. E., S. S. Soliman, X. Li, and D. Bilder. 2011. ‘Altered modes of stem cell division drive adaptive intestinal growth’, Cell, 147: 603–14.

Ohlstein, B., and A. Spradling. 2006. ‘The adult Drosophila posterior midgut is maintained by pluripotent stem cells’, Nature, 439: 470–4.

Oshima, H., M. Nakayama, T. S. Han, K. Naoi, X. Ju, Y. Maeda, S. Robine, K. Tsuchiya, T. Sato, H. Sato, M. M. Taketo, and M. Oshima. 2015. ‘Suppressing TGFbeta signaling in regenerating epithelia in an inflammatory microenvironment is sufficient to cause invasive intestinal cancer’, Cancer Res, 75: 766–76.

Parker, L., J. E. Ellis, M. Q. Nguyen, and K. Arora. 2006. ‘The divergent TGF-beta ligand Dawdle utilizes an activin pathway to influence axon guidance in Drosophila’, Development, 133: 4981–91.

Pauklin, S., and L. Vallier. 2015. ‘Activin/Nodal signalling in stem cells’, Development, 142: 607–19.

Peterson, A. J., and M. B. O’Connor. 2013. ‘Activin receptor inhibition by Smad2 regulates Drosophila wing disc patterning through BMP-response elements’, Development, 140: 649–59.

Reiff, T., J. Jacobson, P. Cognigni, Z. Antonello, E. Ballesta, K. J. Tan, J. Y. Yew, M. Dominguez, and I. Miguel-Aliaga. 2015. ‘Endocrine remodelling of the adult intestine sustains reproduction in Drosophila’, Elife, 4: e06930.

Richmond, C. A., M. S. Shah, L. T. Deary, D. C. Trotier, H. Thomas, D. M. Ambruzs, L. Jiang, B. B. Whiles, H. D. Rickner, R. K. Montgomery, A. Tovaglieri, D. L. Carlone, and D. T. Breault. 2015. ‘Dormant Intestinal Stem Cells Are Regulated by PTEN and Nutritional Status’, Cell Rep, 13: 2403–11.

Rossi, A. M., and C. Desplan. 2020. ‘Extrinsic activin signaling cooperates with an intrinsic temporal program to increase mushroom body neuronal diversity’, Elife, 9.

Secor, S. M., E. D. Stein, and J. Diamond. 1994. ‘Rapid upregulation of snake intestine in response to feeding: a new model of intestinal adaptation’, Am J Physiol, 266: G695–705.

Song, W., D. Cheng, S. Hong, B. Sappe, Y. Hu, N. Wei, C. Zhu, M. B. O’Connor, P. Pissios, and N. Perrimon. 2017. ‘Midgut-Derived Activin Regulates Glucagon-like Action in the Fat Body and Glycemic Control’, Cell Metab, 25: 386–99.

Song, W., A. C. Ghosh, D. Cheng, and N. Perrimon. 2018. ‘Endocrine Regulation of Energy Balance by Drosophila TGF-beta/Activins’, Bioessays, 40: e1800044.

Tian, A., B. Wang, and J. Jiang. 2017. ‘Injury-stimulated and self-restrained BMP signaling dynamically regulates stem cell pool size during Drosophila midgut regeneration’, Proc Natl Acad Sci U S A, 114: E2699–E708.

Ting, C. Y., P. G. McQueen, N. Pandya, T. Y. Lin, M. Yang, O. V. Reddy, M. B. O’Connor, M. McAuliffe, and C. H. Lee. 2014. ‘Photoreceptor-derived activin promotes dendritic termination and restricts the receptive fields of first-order interneurons in Drosophila’, Neuron, 81: 830–46.

Topczewska, J. M., L. M. Postovit, N. V. Margaryan, A. Sam, A. R. Hess, W. W. Wheaton, B. J. Nickoloff, J. Topczewski, and M. J. Hendrix. 2006. ‘Embryonic and tumorigenic pathways converge via Nodal signaling: role in melanoma aggressiveness’, Nat Med, 12: 925–32.

Upadhyay, A., A. J. Peterson, M. J. Kim, and M. B. O’Connor. 2020. ‘Muscle-derived Myoglianin regulates Drosophila imaginal disc growth’, Elife, 9.

Wang, S., and Y. G. Chen. 2018. ‘BMP signaling in homeostasis, transformation and inflammatory response of intestinal epithelium’, Sci China Life Sci, 61: 800–07.

Wells, B. S., D. Pistillo, E. Barnhart, and C. Desplan. 2017. ‘Parallel Activin and BMP signaling coordinates R7/R8 photoreceptor subtype pairing in the stochastic Drosophila retina’, Elife, 6.

Yamada, Y., H. Mashima, T. Sakai, T. Matsuhashi, M. Jin, and H. Ohnishi. 2013. ‘Functional roles of TGF-beta1 in intestinal epithelial cells through Smad-dependent and non-Smad pathways’, Dig Dis Sci, 58: 1207–17.

Yilmaz, O. H., P. Katajisto, D. W. Lamming, Y. Gultekin, K. E. Bauer-Rowe, S. Sengupta, K. Birsoy, A. Dursun, V. O. Yilmaz, M. Selig, G. P. Nielsen, M. Mino-Kenudson, L. R. Zukerberg, A. K. Bhan, V. Deshpande, and D. M. Sabatini. 2012. ‘mTORC1 in the Paneth cell niche couples intestinal stem-cell function to calorie intake’, Nature, 486: 490–5.

Yu, X. M., I. Gutman, T. J. Mosca, T. Iram, E. Ozkan, K. C. Garcia, L. Luo, and O. Schuldiner. 2013. ‘Plum, an immunoglobulin superfamily protein, regulates axon pruning by facilitating TGF-beta signaling’, Neuron, 78: 456–68.

Zeng, X., and S. X. Hou. 2015. ‘Enteroendocrine cells are generated from stem cells through a distinct progenitor in the adult Drosophila posterior midgut’, Development, 142: 644–53.

Zhai, Z., J. P. Boquete, and B. Lemaitre. 2017. ‘A genetic framework controlling the differentiation of intestinal stem cells during regeneration in Drosophila’, PLoS Genet, 13: e1006854.

Zheng, X., C. T. Zugates, Z. Lu, L. Shi, J. M. Bai, and T. Lee. 2006. ‘Baboon/dSmad2 TGF-beta signaling is required during late larval stage for development of adult-specific neurons’, EMBO J, 25: 615–27.

Zhou, J., S. Florescu, A. L. Boettcher, L. Luo, D. Dutta, G. Kerr, Y. Cai, B. A. Edgar, and M. Boutros. 2015. ‘Dpp/Gbb signaling is required for normal intestinal regeneration during infection’, Dev Biol, 399: 189–203.

Zhu, C. C., J. Q. Boone, P. A. Jensen, S. Hanna, L. Podemski, J. Locke, C. Q. Doe, and M. B. O’Connor. 2008. ‘Drosophila Activin-and the Activin-like product Dawdle function redundantly to regulate proliferation in the larval brain’, Development, 135: 513–21.

